# SVCollector: Optimized sample selection for validating and long-read resequencing of structural variants

**DOI:** 10.1101/342386

**Authors:** Fritz J. Sedlazeck, Zachary Lemmon, Sebastian Soyk, William J. Salerno, Zachary Lippman, Michael C. Schatz

## Abstract

**Summary:** Structural Variations (SVs) are increasingly recognized for their importance in genomics. Short-read sequencing is the most widely-used approach for genotyping large numbers of samples for SVs but suffers from relatively poor accuracy. Here we present SVCollector, an open-source method that optimally selects samples to maximize variant discovery and validation using long read resequencing or PCR-based validation. SVCollector has two modes: selecting those samples that are individually the most diverse or those that collectively capture the largest number of variations.

**Availability:** https://github.com/fritzsedlazeck/SVCollector

**Supplementary information:** Supplementary data are available at *Bioinformatics* online.

## 1 Introduction

In recent years it has become increasingly clear that structural variations (SVs) play a key role in evolution, diseases and many other aspects of biology across all organisms (Lupski, 2015; Sudmant, et al., 2015). Short-read sequencing is the most widely-used approach for identifying SVs, although suffers from limited accuracy (Chaisson, et al., 2015; Sedlazeck, et al., 2018). Long reads, such as from PacBio and Oxford Nanopore, provide greater sensitivity and lower false discovery rates, but their higher costs hinder widespread application in large sequencing studies. Another related problem with large cohorts is how to efficiently validate a large number of SVs from the short read-based calls. Traditional methods such as PCR/Sanger sequencing are costly and labor intensive, necessitating careful consideration of variants and samples to validate for further study.

Here we present SVCollector, an open source method (MIT license) to automatically rank and select samples based on variants that are shared within a large population. This allows for both a more cost-efficient way to validate common SVs, along with an optimized approach to discover SVs that were initially missed by short-read sequencing.

## 2 Materials and methods

SVCollector is implemented in C++, and computes a ranked list and diagnostic plots of the samples listed in multi-sample VCF file. It uses an iterative approach to minimize the memory footprint, and requires < 2MB of RAM even when ranking thousands of samples with tens of thousands of variants each. In the first iteration, it parses the VCF file, counts the total number of variants, and generates a temporary file storing the sample ids associated with each SV. For subsequent iterations, it reads the temporary file and deletes SVs that were present in the previously selected sample.

SVCollector has two major ranking modes: *topN*, and *greedy*. For the *topN* mode, it picks samples with the largest number of SVs irrespective if the SVs are shared with other samples. For the *greedy* mode, it finds a set of samples that collectively contain the largest number of distinct variants. Solving this exactly is computationally intractable as it is a version of the well-known NP-hard set cover problem. Consequently, it uses a greedy approximation that starts with the sample with the largest number of variants, and then iteratively picks the sample containing the largest number variants not yet included in the set. It also has a *random* mode that mimics an arbitrary selection process, and is helpful for evaluating the diversity of the topN or greedy approaches. For each mode, SVCollector reports the rank, sample name, its unique contribution of SVs, the cumulative sum of SVs up to the chosen sample, and the cumulative percentage compared to the total number of SVs in the input VCF file.

## 3 Results and Discussion

We assessed the results of SVCollector based on simulated data (**Supplementary Note 1**) and three large short-read sequencing projects involving 354 to 4,242 samples each. For each cohort, we focused on picking the top 100 most diverse samples identified by SVCollector, as compared to a random set. The *random* selection was run 100 times per cohort, and we report the averaged percent of SVs identified. For all cohorts, the runtime and memory requirements were minimal. For example, for the 1000 Genomes VCF file of 2,504 samples over 66,555 distinct SVs (Sudmant, et al., 2015), SVCollector computed the top 100 samples in 67 seconds using 1.7MB memory. Each of the modes had a similar runtime.

### Assessment of wild and domesticated tomatoes

We first ranked 354 samples from a tomato core collection sequenced with short-reads to an average of 30x coverage in the context of GWAS studies on plant domestication and breeding (Tomato Genome Sequencing, et al., 2014; Zhu, et al., 2018) (**Supplementary Note 2**). We identified 201,201 SVs using SURVIVOR (Jeffares, et al., 2017) across all samples. **Figure 1A** shows the results over the top 100 ranked samples over all 3 modes (**Supplementary Table 1**). Among the top 10, the *greedy* approach was able to capture the most SVs (45.40%) followed by the *topN* (41.47%) and *random* (11.80%) approaches. When extending to the top 100, the *greedy* approach again captured more (90.92%) than the *topN* (88.18%) approach followed by the *random* sample selection (49.89%). This difference illustrates the amount of shared vs. singleton SVs across the samples in this cohort (**Supplementary Figure 3**).

**Figure 1:**
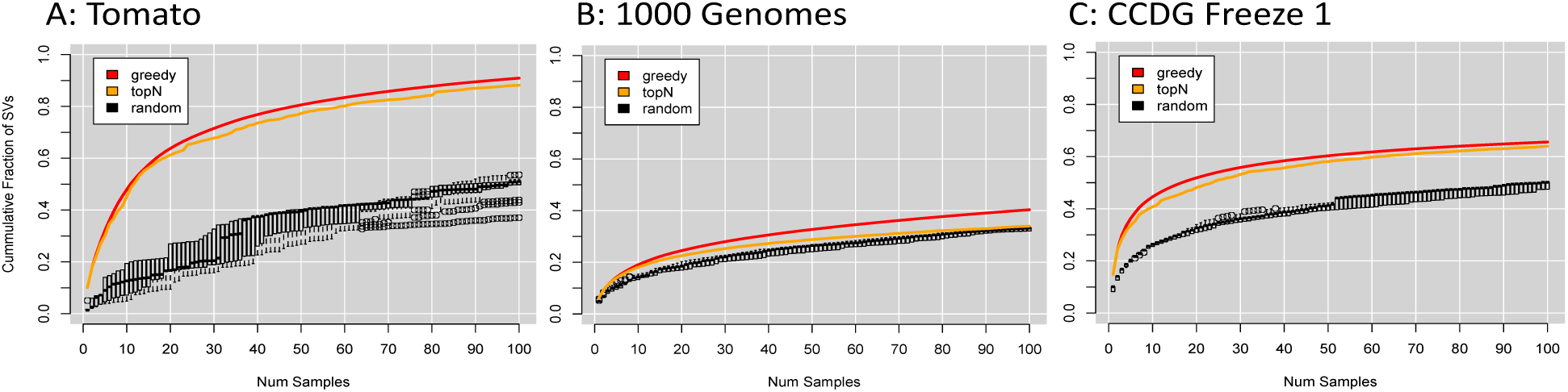
SVCollector sample selection evaluation based on greedy (red), top100 (orange) and random (black) selections.

### Assessment of human genomes

Next, we assessed SVCollector based on 2,504 human genomes from the 1000 Genomes Project (Sudmant, et al., 2015) **Figure 1B** shows the results and **Supplementary Note 3 and Table 3** list the details. The *greedy* approach (19.04%) slightly outperformed the *topN* approach (18.05%) when investigating the first 10 ranked samples. However, when extending the selection to 100 genomes, the *greedy* (40.33%) approach more substantially outperforms the *topN* (33.99%) and *random* (32.81%) approaches. Next, we investigated the sampling across the population groups in the collection (**Supplementary Table 4**). Notably, the *topN* approach selected 99 African and 1 American samples, while the *greedy* approach covered all 8 populations representing 25 of the 26 subpopulations, excluding only GBR, and contained 60 African, 16 South Asian, 14 East Asian, 6 European and 4 American samples.

Finally, we assessed the performance of SVCollector based on 4,424 human genomes from the Center for Common Disease Genetics (CCDG) freeze 1 dataset composed of 425,500 SVs identified with SURVIVOR (Manuscript in preparation). **Figure 1C** shows the results and **Supplementary Note 4 and Table 5** lists the details. As in the other datasets, the greedy approach over the top 10 samples captured the most SVs compared to the *topN* and *random* approach with 43.39%, 39.92% and 25.26%, respectively. Extending the selection to 100 samples captured 65.58%, 63.98% and 48.86% for *greedy*, *topN* and *random* approach, respectively.

## 4 Conclusion

SVCollector is a fast and powerful method to quantitatively select samples for long-read resequencing or PCR-validation based on their genomic variations shared in the population. Both the *topN* and *greedy* modes can substantially outperform the commonly used *random* selection. This in turn will translate to a substantial cost savings for resequencing and validation studies. Interestingly, in the 1000 Genomes dataset, the topN approach was essentially equivalent to picking a random subset of genomes, and both could be outperformed by picking an optimal set with the *greedy* approach. We also noted that more than 100 samples were needed to capture 50% of the diversity of the 1000 Genomes population, but less than 20 were needed for the other two populations, suggesting more constrained diversity. In the human datasets, female samples often contributed more SVs than male samples because of the extra heterozygous SVs on the X chromosome. Depending on the application, researchers may want to filter out the sex chromosomes prior to analysis. Relatedly, SVCollector can also operate on multi-sample VCFs with SVs and SNPs, and we find these approaches can produce different rankings of samples depending on the needs of the project (**Supplementary Note 2**).

## Funding

National Science Foundation DBI-1350041 and IOS-1732253, National Institutes of Health R01-HG006677 and UM1-HG008898.

